# Increasing the accuracy of exchange parameters reporting on slow dynamics by performing CEST experiments with high *B*_1_ fields

**DOI:** 10.1101/2024.04.02.587659

**Authors:** Nihar Pradeep Khandave, D Flemming Hansen, Pramodh Vallurupalli

## Abstract

Over the last decade chemical exchange saturation transfer (CEST) NMR methods have emerged as powerful tools to characterize biomolecular conformational dynamics occurring between a visible major state and ‘invisible’ minor states. The ability of the CEST experiment to detect these minor states, and provide precise exchange parameters, hinges on using appropriate *B*_1_ field strengths during the saturation period. Typically, a pair of *B*_1_ fields with *ω*_1_ (= 2*πB*_1_) values around the exchange rate *k*_ex_ are chosen. Here we show that the transverse relaxation rate of the minor state resonance (*R*_2,*B*_) also plays a crucial role in determining the *B*_1_ fields that lead to the most informative datasets. Using 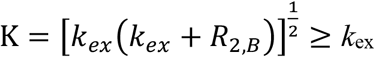, to guide the choice of *B*_1_, instead of *k*_ex_, leads to data wherefrom substantially more accurate exchange parameters can be derived. The need for higher *B*_1_ fields, guided by K, is demonstrated by studying the conformational exchange in two mutants of the 71 residue FF domain with *k*_ex_ ∼11 s^-1^ and ∼72 s^-1^, respectively. In both cases analysis of CEST datasets recorded using *B*_1_ field values guided by *k*_*ex*_ lead to imprecise exchange parameters, whereas using *B*_1_ values guided by K resulted in precise site-specific exchange parameters. The conclusions presented here will be valuable while using CEST to study slow processes at sites with large intrinsic relaxation rates, including carbonyl sites in small to medium sized proteins, amide ^15^N sites in large proteins and when the minor state dips are broadened due to exchange among the minor states.

## Introduction

Protein molecules are dynamic entities that at ambient temperature sample various conformational states with differing populations and lifetimes (1, 2). In addition to understanding dynamical processes, such as protein folding/misfolding and aggregation, a knowledge of protein conformational dynamics is often necessary to understand protein function, allostery etc. (2-6). Hence, over the last few decades different classes of NMR experiments have been developed to study protein conformational dynamics occurring on the μs to second time-scale (4, 7-9), including *R*_1,*ρ*_ (10, 11), CPMG (12, 13), CEST (14) and DEST (15). These experiments can detect sparsely populated conformational states that are ‘invisible’ in regular NMR spectra. In all these experiments, the spins are manipulated by pulses whereafter the ‘visible’ major state magnetization is detected and used to reconstruct the spectrum of the ‘invisible’ minor state, which in favorable cases can be used to determine the structures of the minor states (16-20). CEST experiments, originally devised to study slow exchange between visible states (21), are now routinely used to study protein and nucleic acid conformational exchange between a visible major state and invisible minor state(s) occurring over a wide range of time-scales (22-24). CEST methods have been developed to characterize the exchange at various backbone and side-chain sites (25-31) and have been used to study various processes, including protein folding (24, 32), ligand binding (33, 34) and several other processes involving protein and nucleic acid conformational fluctuations (35-37).

In a typical CEST experiment longitudinal magnetization arising from the nucleus of interest is irradiated with a weak *B*_1_ (∼5 to ∼300 Hz) field for a period *T*_EX_ of ∼0.25 to ∼0.6 s termed the exchange delay, following which the intensity of the visible major-state peak is quantified as a function of the offset at which the *B*_1_ irradiation is applied. When the system of interest consists of a major state, A, in slow exchange with a minor state, B, that is 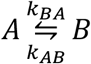, a plot of the normalized intensity 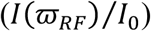 *versus* the offset 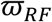 (ppm) at which the *B*_1_ field is applied will have two dips. These two dips consist of one at the chemical shift (ppm) of the major state 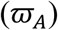 and more importantly one at the chemical shift of the minor state 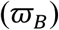, which allows one to detect sparsely populated states with fractional populations as low as ∼0.5%. *I*_0_ is the intensity of the major state in the absence of the *T*_*EX*_ delay. The size and width of the minor state dip (largely) depends on the exchange rate (*k*_*ex*_ = *k*_*AB*_ + *k*_*BA*_), the fractional population of the minor state (*p*_*B*_ = *k*_*AB*_/*k*_*ex*_), the minor state transverse relaxation rate (*R*_2,*A*_) and the value of *B*_1_. The exchange parameters (*k*_*ex*_, *p*_*B*_), the major and minor state chemical shifts, the major (*R*_2,*A*_) and the minor-state transverse relaxation rates (*R*_2,*B*_), as well as the major state longitudinal relaxation rate (*R*_1,*A*_) can all be extracted from the analysis of a pair of CEST profiles recorded with different (suitably chosen) *B*_1_ values (14). For two-state slow exchange (*k*_*ex*_/|Δ*ω*_*AB*_| < 1) processes considered here, CEST profiles are typically recorded with *ω*_1_ (rad/s; = 2 *B*_*1*_) values guided by *k*_*ex*_, that is one *ω*_1_ less than *k*_*ex*_ in the 0.5*k*_*ex*_ to 0.8*k*_*ex*_ range and one higher than *k*_*ex*_ in the 1.5*k*_*ex*_ to 1.8*k*_*ex*_ range. Here Δ*ω*_*AB*_ = *ω*_*B*_ − *ω*_*A*_, where *ω*_*A*_ and *ω*_*B*_ are the resonance frequencies (rad/s) of the nucleus of interest in the major and minor states respectively.

The small 71 residue four helix bundle FF domain (38) from human HYPA/FBP11 has served as a model system to understand protein conformational dynamics and folding (39-44). Whilst characterizing the conformational dynamics of the A17G S56P FF domain using methyl ^13^C CEST experiments, we found that precise site-specific exchange parameters could not be obtained from the analysis of two CEST datasets recorded with *ω*_1_ values lower and higher than *k*_*ex*_ (∼11 s^-1^) as described above. We discovered that this problem occurs when the transverse relaxation rate of the minor state, *R*_2,*B*_, is greater than *k*_*ex*_ and that accurate exchange parameters can be obtained by recording additional CEST datasets with higher *B*_1_ values. We rationalize the benefit of the larger *B*_1_ for deriving accurate exchange parameters by inspecting the equations that govern the shape of the minor state dip in CEST profiles. We conclude that the choice of *B*_1_ values should be informed by 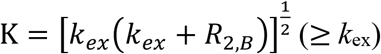 rather than *k*_*ex*_. The efficacy of this strategy is further demonstrated by characterizing the two-state folding reaction (*k*_*ex*_ ∼ 72 s^-1^) of the A39G FF domain in which the apparent transverse relaxation rates of several residues in the unfolded (U) state are greater than 140 s^-1^. In line with the analysis presented here, choosing *B*_1_ values guided by K rather than *k*_*ex*_ allows for the accurate determination of the exchange parameters.

## Materials and Methods

### Protein Samples

The A17G S56P FF sample contained of ∼4 mM U-[^2^H,^15^N], Ileδ1-[^13^CH_3_], Leu,Val-[^13^CH_3_,^12^CD_3_] labelled protein dissolved in 50 mM acetate, 100 mM NaCl, 30% [^2^H]-glucose, pH 5.7, 100% D_2_O buffer. The A39G FF sample contained ∼4 mM U-[^15^N] labelled protein dissolved in 50 mM acetate, 100 mM NaCl, pH 5.7, 10% D_2_O buffer. Proteins were overexpressed in *E coli* BL21(DE3) cells transformed with the appropriate plasmids grown in suitable M9 media (45, 46) and purified as described previously (24, 47, 48).

### NMR Spectroscopy

Methyl ^13^C CEST profiles (A17G S56P FF sample, 7.5 °C) were recorded on a 700 MHz (16.4 T) Bruker Avance III HD spectrometer equipped with a cryogenically cooled triple resonance probe. To accelerate data acquisition, ^13^C methyl CEST data was acquired using the DANTE-CEST (D-CEST) sequence (49, 50) that uses the DANTE sequence (51, 52) for RF irradiation during the *T*_*EX*_ period. Amide ^15^N CEST profiles (A39G FF sample, 2.5 °C) were recorded on a 500 MHz (11.7 T) Bruker NEO spectrometer equipped with a room temperature triple resonance probe using the standard amide ^15^N CEST sequence (25). During the *T*_*EX*_ delay of both the ^13^C and ^15^N CEST experiments ^1^H decoupling was carried out using the 90_x_240_y_90_x_ composite pulse (53) effectively reducing the nucleus of interest (methyl ^13^C or the amide ^15^N) to an isolated spin ½ spin system (25). *B*_*1*_ fields were calibrated using the nutation method (54). The methyl ^1^H-^13^C correlation maps were recorded with 24 complex points (sweep width: 14 ppm) in the indirect (^13^C) dimension while the amide ^1^H-^15^N correlation maps were recorded with 24 complex points (sweep width: 16.9 ppm) in the ^15^N dimension. Methyl ^13^C CEST data were acquired using 16 scans, whereas 4 scans were used to record the amide ^15^N CEST data. Additional details are provided in Table S1.

### Data Analysis

The NMRPipe package (55) was used to process the NMR data, Sparky (56, 57) was used to visualize and label the spectra while the program PINT (58) was used to obtain peak intensities from the spectra. Uncertainties in the peak intensities were estimated based on the scatter in the flat part of the CEST intensity profiles (23). The software package *ChemEx* (59) that numerically integrates (60) the Bloch-McConnell equations (61) was used to both obtain the best fit exchange parameters from the experimental (or synthetic) data and to generate the synthetic CEST profiles (Fig. 2 & S2). The two-state fitting parameters included the major and minor state chemical shifts and transverse relaxation rates, the major state longitudinal relaxation rate (*R*_1,*A*_) and the exchange rate and the minor state population. While fitting data from multiple sites to a global two-state process the exchange rate and minor state population were assumed to be the same for all sites. In all the data analysis the longitudinal relaxation rate was assumed to be the same for both states. Unless mentioned uncertainties in the best fit exchange parameters were estimated using a standard Monte Carlo procedure that consisted of 250 trials (62, 63).

## Results and Discussion

### The choice of optimal *B*_*1*_ fields can depend on the minor state transverse relaxation rate in addition to *k*_*ex*_

The A17G S56P FF domain exchanges between the folded state (F) and an alternate conformer (I). The (ILV) methyl ^1^H-^13^C correlation map of U-[^2^H,^15^N], Ileδ1-[^13^CH_3_], Leu,Val-[^13^CH_3_,^12^CD_3_] A17G S56P FF is well resolved at 7.5 °C (Fig. 1a) and a minor state dip is clearly visible in the methyl ^13^C CEST profiles (Fig. 1b) from six sites (V30γ2, I43δ1, I44δ1, L52δ2, L55δ1 & L55δ2). Unlike CPMG experiments where precise exchange parameters are often obtained by a global analysis of data recorded from several sites at multiple *B*_*0*_ fields (64), precise two and even three-state (slow) exchange parameters can be obtained on a per site basis by analyzing CEST data recorded at a single *B*_*0*_ field, but with multiple *B*_1_ fields instead, allowing one to identify global exchange processes (25, 32). Since the exchange rate was expected to be approximately 10 s^-1^, we initially chose *B*_1_ fields of 1.5 and 3.4 Hz (*ω*_1_ of 9.4 rad/s and 21.4 rad/s). However, when the ^13^C CEST profiles (*B*_*1*_ = 1.5 & 3.4 Hz) from each of the six sites were analyzed independently to obtain site-specific exchange rates *k*_*ex*_ and minor-state fractional populations, *p*_*I*_ (Fig. 1c,d), the extracted two-state exchange parameters were poorly defined. This was particularly the case for the exchange rates, *k*_*ex*_, as shown in Fig. 1c, where the *k*_*ex*_ and *p*_*I*_ values obtained for each of the six sites from a Monte Carlo procedure with 250 trials are plotted (grey circles) and in Fig. 1d where the distributions of the site specific *k*_*ex*_ and *p*_*I*_ values are plotted. The best fit *k*_*ex*_ values range from 6.4 to 51.7 s^-1^, whereas the best fit *p*_*I*_ values range from 5.4 to 12.9 % across the six residues (Table S2). Analysis of ^13^C CEST profiles recorded with *B*_*1*_ = 9.8 Hz in addition to the ones recorded with *B*_*1*_ = 1.5 and 3.4 Hz resulted in more precise *k*_*ex*_ and *p*_*I*_ values (Fig. 1c, blue pluses; compare *k*_*ex*_ distributions in Fig. 1d and 1e) with site-specific best fit *k*_*ex*_ values now varying from 10.1 to 12.4 s^-1^ and best fit *p*_1_ values varying from 8.5 to 10.7 % across the six residues (Table S2). The fact that the analysis of CEST profiles from each of the six sites resulted in very similar exchange parameters (Fig. 1c,e) strongly suggests that they are all reporting on the same global exchange process and a global analysis of the ^13^C CEST data (*B*_*1*_ = 1.5, 3.4 & 9.8 Hz) from all six sites resulted in good quality fits 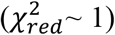 and *k*_*ex*_= 11.2 ± 0.5 s^-1^ and *p*_*I*_= 9.5 ± 0.3 %. Addition of the *B*_*1*_ = 9.8 Hz ^13^C CEST dataset into the analysis procedure leads to a narrower minimum especially for *k*_*ex*_ even in the 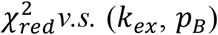 plots (Fig. 1f *v*.*s*. 1g) obtained from a global analysis of CEST data from all six sites.

**Fig. 1.**
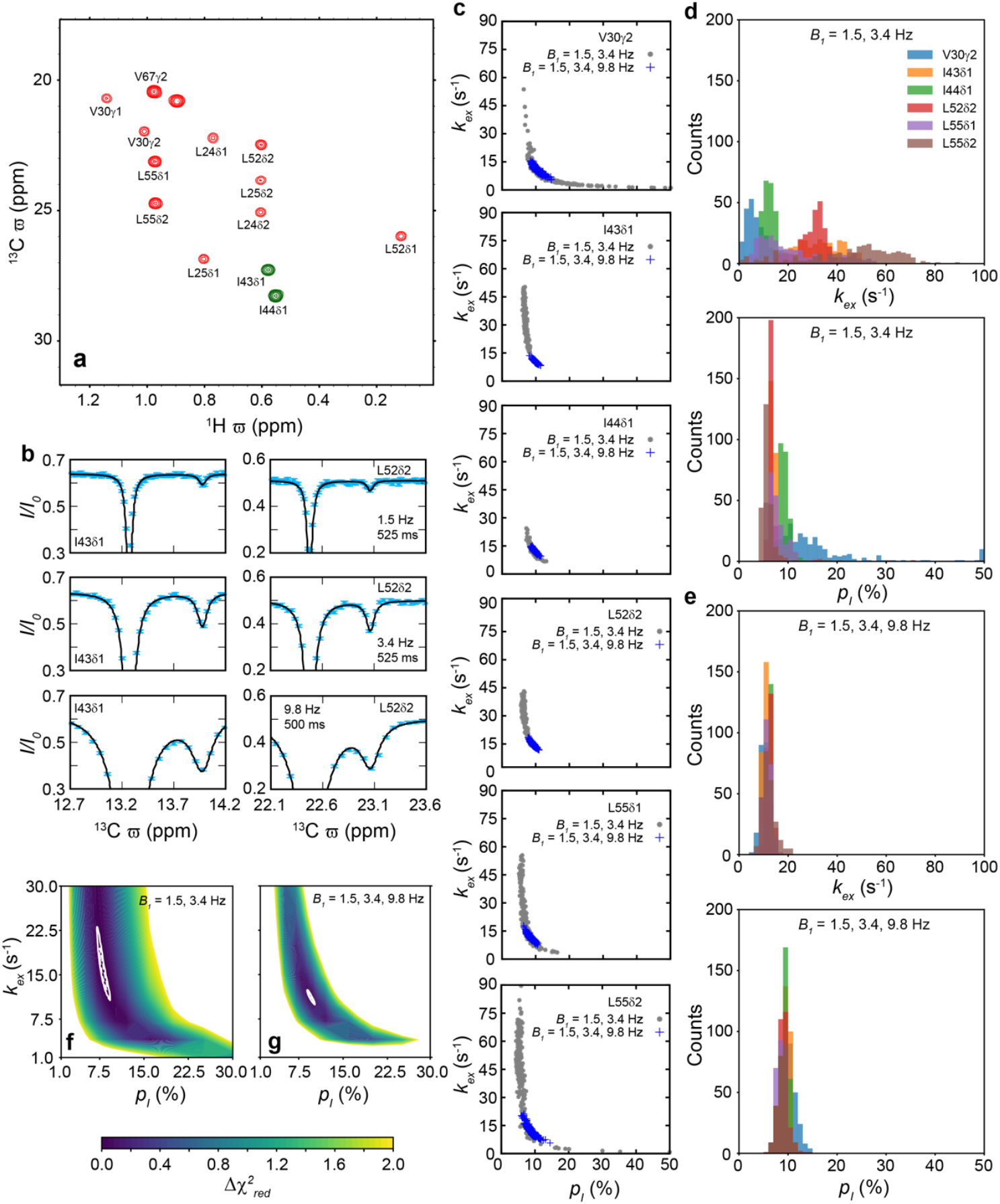
High *B*_*1*_ fields lead to precise exchange parameters for the A17G S56P FF *F ⇌ I* reaction (*k*_*ex*_ ∼11.2 s^-1^). (a) Methyl ^13^C-^1^H correlation map of the U-[^2^H,^15^N],Ileδ1-[^13^CH_3_],Leu,Val-[^13^CH_3_,^12^CD_3_] A17G S56P FF (16.4 T, 7.5 °C). Peaks are labelled according to the site from which they arise. Green peaks are aliased in the ^13^C dimension. (b) Representative methyl ^13^C CEST profiles (*B*_*1*_ and *T*_*EX*_ indicated) clearly show the presence of a minor state dip. Cyan circles are used to represent the experimental data while the black line is drawn using the global best fit parameters (*k*_*ex*_ = 11.2 s^-1^, *p*_1_ = 9.52%; Table S2). (c) Scatter plots showing the distribution of *k*_*ex*_ and *p*_1_ values obtained using a Monte Carlo procedure with 250 trials. Analysis was carried out separately at each site using two different combinations of CEST datasets: *B*_*1*_ values of 1.5 and 3.4 Hz (grey circles) and *B*_*1*_ values of 1.5, 3.4 and 9.8 Hz (blue pluses). (d,e) Histograms showing the distribution of site specific *k*_*ex*_ and *p*_1_ values from (c). 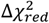 vs (*k*_*ex*_, *p*_1_) plots obtained from a global analysis of the methyl ^13^C CEST 1.5 and 3.4 Hz (f) and 1.5, 3.4 and 9.8 Hz (g) datasets.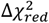 is difference between 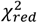 and the minimum (best fit) value of 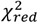 (lowest value of 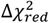 is 0). 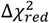 values above 2 are in white. In f and g contours corresponding to the 68 and 95% confidence intervals of *k*_*ex*_ and *p*_*I*_ based on 10,000 Monte Carlo trials are also shown using dashed and solid white lines respectively.

For a *k*_*ex*_ value of 11.2 s^-1^, *B*_*1*_ values of 1.5 and 3.4 Hz correspond to *ω*_1_/*k*_*ex*_ values of 0.8 and 1.9 respectively and this choice of CEST datasets should have sufficed (14) to obtain precise estimates of the exchange parameters unlike what was observed (Fig. 1c,d). To resolve this conundrum, we noted that the fitted *R*_2,*I*_ values (∼20 to ∼70 s^-1^) are all higher than *k*_*ex*_ (∼11 s^-1^) for the six sites (Table S2) and we therefore investigated more generally the effect of the minor state transverse relaxation rate on the size of the minor state dip in CEST profiles.

For a two-state (*A* ⇌ *B*) reaction (*k*_*ex*_ = 15 s^-1^, *p*_*B*_ = 7.5%) the calculated intensity of the minor (*B*) state dip is plotted in Fig. 2a as a function of *ω*_1_ for different *R*_2,*B*_ values while the inset shows the minor state dip for various *R*_2,*B*_ values when *ω*_1_ = 15 rad/s (= *k*_*ex*_). It is clear that when *ω*_1_ is fixed to 15 rad/s, the size of minor state dip decreases as the *R*_2,*B*_ values increase (inset Fig. 2a). For example, the minor dip that is prominent when *R*_2,*B*_ = 5 s^-1^ (black curve Fig. 2a inset) becomes essentially invisible when *R*_”,*B*_ is increased to 125 s^-1^ (cyan curve Fig. 2A inset). A physical explanation is that as *R*_2,*B*_ increases, the *B*_*1*_ field (analogous to *B*_*0*_ under free pression) becomes less effective at inducing a relative phase change between the magnetization exchanging between states B and A. When *R*_2,*B*_ = 5 s^-1^, the intensity of the minor state dip has a distinctive dependence on *ω*_1_ as *ω*_1_/*k*_*ex*_ is varied between ∼0.5 and ∼2 (black curve in Fig. 2a). However, for large transverse relaxation rates in the minor state, *e*.*g. R*_2,*B*_ = 125 s^-1^, the size of the minor state dip is small and its intensity changes to a lesser degree when *ω*_1_/*k*_*ex*_ is varied from 0.5 to 2 (cyan curve in Fig. 2a). Thus, *R*_2,*B*_ influences the size of the minor state dips (65, 66) and when *R*_2,*B*_ is substantially larger than *k*_*ex*_, CEST datasets recorded with *ω*_1_ values much larger than *k*_*ex*_ will be required to see the minor state dip clearly and to obtain accurate exchange parameters (Fig. 2a).

**Fig. 2.**
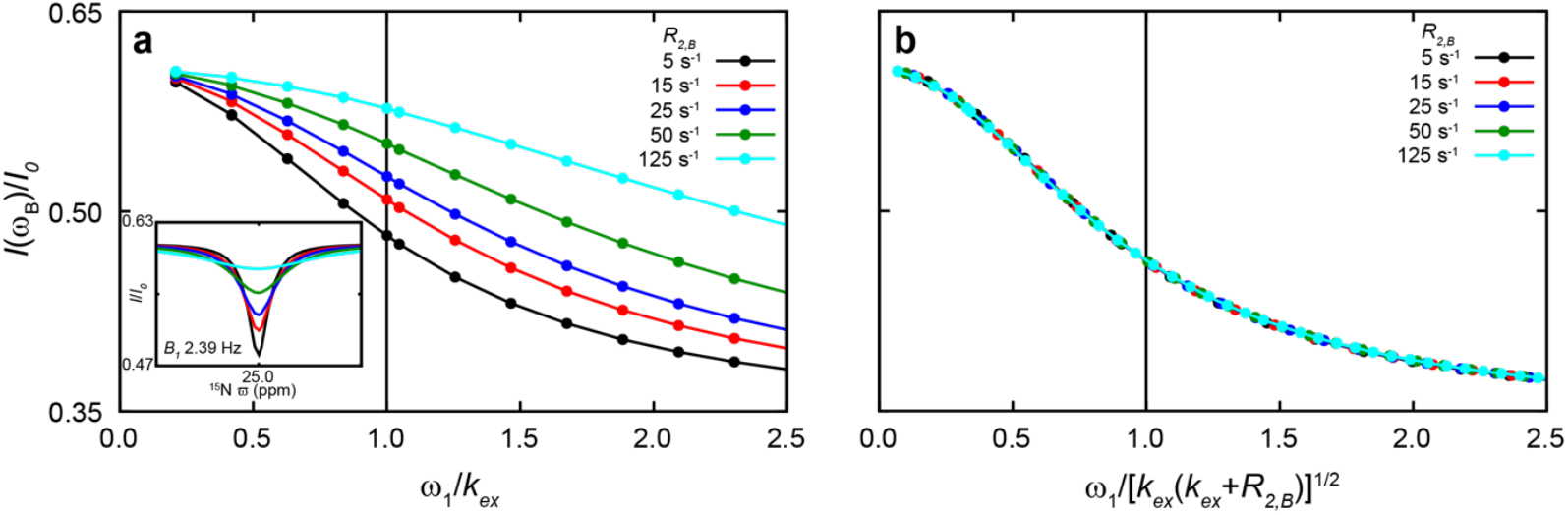
The size of the minor (B) state dip depends on *R*_2,*B*_. Plot of the normalized minor state dip intensity (*I*(*ω*_*B*_)/*I*_0_) as a function of *ω*_1_/*k*_*ex*_ (a) and 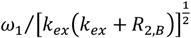 (b) for different *R*_2,*B*_ values. The inset in (a) shows the CEST profile (*B*_1_ = *k*_*ex*_/2*π* = 2.39 Hz) around 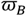 for different *R*_2,*B*_ values. Calculations were performed for a two-state slow exchange reaction (*k*_*ex*_/*Δω*_*AB*_ ∼0) with *k*_*ex*_ = 15 s^-1^, *p*_*B*_ = 7.5 %, *R*_1,*A*_ = 1 s^-1^, *R*_1,*B*_ = 1 s^-1^, *R*_2,*A*_ = 5 s^-1^,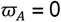ppm, 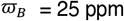 (^15^N, 16.4 T) and *T*_*EX*_ = 0.5 s.

### CEST datasets with *B*_*1*_ fields much larger than *k*_*ex*_/2*π* are required to study exchange when *R*_2,*B*_ is comparable to or larger than *k*_*ex*_

To understand the differing shapes of the *I*(*ω*_*B*_)/*I*_0_ *v*.*s. ω*_1_/*k*_*ex*_ plots in Fig. 2a, we consider a spin ½ nucleus undergoing conformational exchange in the slow exchange regime with 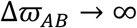. Based on the equivalence between CEST and *R*_1,*ρ*)_ experiments (35, 65, 67) the decay of the ground state magnetization under weak *B*_*1*_ irradiation can be described by 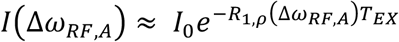 (65, 68) with,

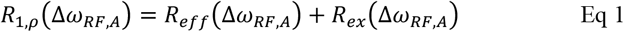

Here Δ*ω*_*RF,i*_ is the difference (rad/s) between the offset at which the *B*_*1*_ irradiation is applied and the resonance frequency of the nucleus in state *i. R*_*eff*_ (Δ*ω*_*RF,A*_) is the effective relaxation rate of the spin under *B*_*1*_ irradiation in the absence of exchange and *R*_*ex*_ (Δ*ω*_*RF,A*_), is the exchange contribution to relaxation. Different expressions have been obtained for *R*_*ex*_ (Δ*ω*_*RF,A*_), (67, 69). Focusing on the minor state and assuming that the longitudinal relaxation rate is 0 s^-1^ the following simple relation (65, 66) for *R*_*ex*_(Δ*ω*_*RF,B*_) applies,

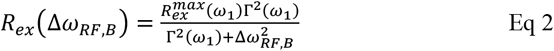

When the RF-irradiation is applied at the offset of the minor state resonance, 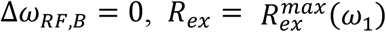, which is the maximum value of *R*_*ex*_ for a given *B*_*1*_. Γ is the half width at the half maximum of *R*_*ex*_, (Fig. S1a). 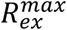 and Γ are given by,

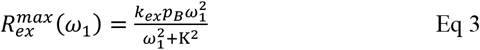

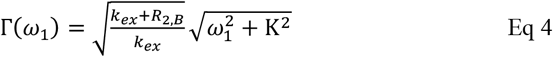

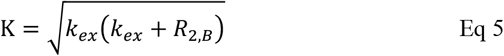

The size of the minor state dip is given by 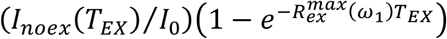, while the shape of the minor state dip (*I*(Δ*ω*_*RF,B*_,/*I*_0_ *v*.*s* Δ_*RF,B*_) is proportional to 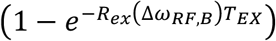, (Fig. S1b). Here *I*_*noex*_(*T*_*EX*_) is the intensity measured at *ω*_*B*_ in the absence of exchange, or equivalently at an offset far from *ω*_*A*_ and *ω*_*B*_ in the presence of exchange, and *I*_*noex*_(*T*_*EX*_) essentially accounts for longitudinal relaxation during *T*_*EX*_. According to Eq 3 the shape of the *I*(*ω*_*B*_)/*I*_0_ *v*.*s ω*_1_/*k*_*ex*_ plots in Fig. 2a is determined by the ratio of *ω*_1_ and K, rather than the ratio of *ω*_1_ and *k*_*ex*_, which means that the curves in Fig. 2a should be identical when *I*(*ω*_*B*_)/*I* is plotted against *ω*_1_/K, as can be seen in Fig. 2b. When *ω*_1_ ≪ K the minor state dip will not be prominent and its size will increase when *ω*_1_ is increased (Fig. 2b, Eq 3, 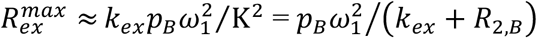) while its width will barely increase when *ω*_1_ is increased (Eq 4; 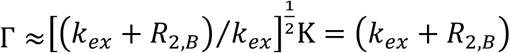). Note that Γ is the half width at the half maximum of *R*_*ex*_(Δ*ω*_*RF,B*_ Eq 3, 4 and Fig. S1, whereas the width of the dip, 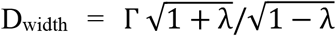, where 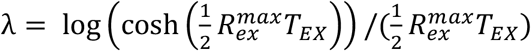, which depends on Γ Provided that the SNR is adequate to see the minor state dip, analysis of CEST profiles recorded with *ω*_1_ ≪ K can lead to reasonable estimates of *p*_*B*_ (compare 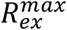 and Γ) but not *k*_*ex*_ as *k*_*ex*_ cannot be separated from *R*_*RF,B*_. On the other hand, when *ω*_1_ ≫ K the minor state dip will be prominent, but its size will be independent of *ω*_1_ (Fig. 2b, Eq 3, 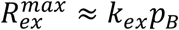) while its width will increase when *ω*_1_ is increased (Eq 4, 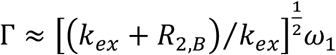). Only the forward rate, *k*_*AB*_ = *k*_*ex*_*p*_*B*_ can be estimated by analyzing of CEST profiles recorded with *ω*_1_ ≫ K. Thus, recording multiple profiles exclusively with *ω*_1_ ≫ K, or exclusively with *ω*_1_ ≪ K, will not provide any additional information and will not meaningfully aid in estimating accurate exchange parameters. The intensity (and width, Eq 4) of the minor state dip shows a distinctive dependence on the value of *ω*_1_, when *ω*_1_ ∼ K (Fig. 2b) making it clear that, in order to derive accurate exchange parameters, CEST datasets should be recorded with *B*_*1*_ values guided by K. For example, it follows from the above discussion that accurate *k*_*ex*_ and *p*_*B*_ values can be obtained from a combined analysis of CEST profiles recorded with *ω*_1_ < K and *ω*_1_ > K because *p*_*B*_ can effectively be derived from CEST profiles recorded with *ω*_1_ < K and *k*_*ex*_*p*_*B*_ can be estimated by analyzing CEST profiles recorded with *ω*_1_ > K. K is larger than *k*_*ex*_ and it begins to deviate significantly from *k*_*ex*_ as the value of *R*_”,*B*_ becomes greater than *k*_*ex*_. The above discussion follows the analysis presented previously (14), except for the fact that the effects of *R*_*RF,B*_ have been explicitly retained here and as expected when *R*_*RF,B*_ ≪ *k*_*ex*_, K ∼ *k*_*ex*_ leading to the previous conclusion that to obtain accurate two-state exchange parameters CEST datasets should be recorded using *B*_*1*_ values informed by *k*_*ex*_. For a global process, if the value of *R*_*RF,B*_ is constant across the molecule *i*.*e*. same K for all sites under investigation then a pair of CEST datasets recorded with *ω*_1_ values in the (0.5-0.8)K and (1.5-1.8)K ranges will suffice to obtain accurate exchange parameters (14). The dataset with *ω*_1_ in the (0.5-0.8)K range will have small and unbroadened minor state dips, whereas the dataset with *ω*_1_ in the (1.5-1.8)K range will have prominent but (*ω*_1_) broadened minor state dips. However, *R*_*RF,B*_ may not be constant throughout the molecule, as in the cases studied here, and in such cases it may not be possible to obtain precise exchange parameters from just two CEST datasets. Hence it will be useful to record an additional CEST dataset with a relatively high *B*_*1*_ so that *ω*_1_/K is greater than ∼1.8 for all sites in the molecule to supplement the datasets recorded with lower *B*_*1*_ values (guided by *k*_*ex*_), where *ω*_1_/K samples some part of the 0.5 to ∼1 region for all residues. A *B*_*1*_ value for which 2*πB*_1_/*k*_*ex*_ ∼4.5 may serve as a starting *B*_*1*_ value for the additional (high *B*_*1*_) dataset as this will result in *ω*_1_/K ∼1.8 even when *R*_*RF,B*_ is relatively high ∼5*k*_*ex*_. If an estimate of *k*_*ex*_ is not available, approximate ranges for *k*_*ex*_ and *R*_*RF,B*_ can be estimated by analyzing preliminary CEST data that preferably contains a dataset recorded with a relatively high *B*_*1*_, for example 50 Hz. K calculated from these estimates can then be used to guide the choice of *B*_*1*_ values to record additional CEST datasets. It should be noted that the minor state *R*_*2*_ values affect the choice of *B*_*1*_ fields used in DEST experiments where resolving the minor state dip is not a concern (9, 15). The validity of the analysis presented above has been confirmed using Monte Carlo simulations (See supporting text and figure S2).

The above analysis can be used to rationalize the previous observation, that precise exchange parameters could not be extracted for the A17G S56P FF domain *F* ⇌ *I* process (*k*_*ex*_ ∼11 s^-1^) by analyzing CEST datasets recorded with *B*_*1*_ = 1.5 (*ω*_1_/*k*_*ex*_ ∼ 0.8) and 3.4 Hz (*ω*_1_/*k*_*ex*_ ∼ 1.9), but accurate exchange parameters could be extracted upon the inclusion of an additional CEST dataset recorded with a relatively high *B*_*1*_ = 9.8 Hz (*ω*_1_/*k*_*ex*_ ∼ 5.5) in the least-squares fit procedure. As mentioned earlier the fitted *R*_2,*I*_ values at various sites varied from ∼20 to ∼70 s^-1^ all of which are substantially higher than *k*_*ex*_. For a *R*_2,*I*_ of 25 s^-1^, K ∼20 s^-1^, and consequently *B*_*1*_ values of 1.5 and 3.4 Hz correspond to *ω*_1_/K values of ∼0.5 and ∼1.1 respectively that are lower than the desired *B*_*1*_ values required to obtain accurate exchange parameters. A *B*_*1*_ value of 9.8 Hz corresponds to *ω*_1_/K of 3.1 when *R*_2,*I*_ is 25 s^-1^ and ∼2 when *R*_2,*I*_ is 70 s^-1^ and therefore including a dataset recorded with *B*_*1*_ = 9.8 Hz in the analysis procedure provides the desired higher *B*_*1*_ dataset.

To further test the above strategy, we have used amide ^15^N CEST experiments to characterize the folding of the A39G FF domain because the minor state dips in the ^15^N CEST profiles are severely broadened due to additional exchange. A39G FF folds from the unfolded state (U) to the native state (F) via two intermediates (I1 and I2) at a rate of ∼70 s^-1^ (3 °C) with *p*_u_ ∼1 %, *p*_*I*1_∼0.3 % and *p*_*I2*_∼0.2 % (24). As U and the folding intermediates I1 and I2 rapidly interconvert among each other on the ∼0.1 to ∼1 ms timescale, the folding reaction can be treated as a two-state exchange reaction between the native state (F) and a state U’. U’ which is a composite of U, I1 and I2 can be described using a combination of the exchange parameters that are used to describe U, I1 and I2 (24). For example, *p*_u′_ ≈ *p*_u_ + *p*_*I*1_ + *p*_*I2*_ and 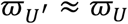 is slightly shifted from 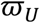 towards 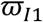 because U and I1 are in fast exchange (24). Exchange between U, I1 and I2 severely broadens several U’ dips and explicit dips arising from the I1 state are not visible in any of the CEST profiles, whereas the CEST profile of only Ser 56 (that is excluded from the present analysis) contains an explicit dip due to the I2 state (24). The amide ^15^N-^1^H correlation map is well resolved (Fig. 3a) and ^15^N CEST profiles were obtained (Fig. 3b) for 58 out 60 (non-proline) ordered (residue 10 to 71) amino acid sites in the molecule. In the discussion that follows we only consider 19 sites with large chemical shift differences 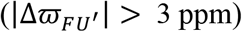. A global two-state exchange model satisfied 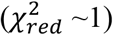 the (*B*_*1*_ = 6.0 & 18.4 Hz) ^15^N CEST data resulting in well-defined exchange parameters with *k*_*ex*_ = 72 ± 3 s^-1^ and *p*_2,U,_′ = 1.39 ± 0.03 % and the CEST derived 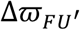 values are in good agreement with the predicted 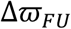 values (RMSD 1.8 ppm, Fig. 3c) confirming that the U state is the dominant state among the states that comprise U’. As mentioned above, the exchange between U, I1 and I2 results in some U’ dips that are severely broadened, as can be seen in Fig. 3b where the minor state dip of I43 is significantly broader than that of L52 and Q68. The distribution of *R*_2,U,_′ values obtained from the global two-state analysis of (*B*_*1*_ = 6.0 & 18.4 Hz) amide ^15^N CEST profiles is plotted in Fig. 3d (Table S3). The *R*_2,U,_′values show a broad distribution with several residues having *R*_2,U,_′ values above 2*k*_*ex*_ (∼140 s^-1^). For the four residues (L52, L55, K66 & Q68; Table S3) with *R*_2,U,_′values less than 50 s^-1^ (K = 94 s^-1^ when *R*_2,U,_′ = 50 s^-1^) single residue fits of ^15^N CEST data (*B*_*1*_ = 6.0 & 18.4 Hz) yielded well defined exchange parameters (Fig. 3e) with *k*_*ex*_ varying from 53 to 73 s^-1^ and *p*_U′_ varying from 1.4 to 1.7 % across the four different residues (Table S3). Including an additional CEST dataset recorded with *B*_*1*_ = 46 Hz in the fitting procedure only has a small effect on the exchange parameters extracted for these residues (Fig. 3f) with *k*_*ex*_ now varying from 57 to 64 s^-1^ and *p*_U′_ now varying from 1.4 to 1.6 % across the different residues. In contrast, for the seven residues (T13, K28, R29, M42, I43, I44 & N45; Table S3) with *R*_2,U′_ values greater than 140 s^-1^ (∼2*k*_*ex*_), analysis of the *B*_*1*_ = 6.0 and 18.4 Hz ^15^N CEST datasets, on a per residue basis, resulted in poorly defined exchange parameters (Fig. 3g) with *k*_*ex*_ varying from 40 to 146 s^-1^ and *p*_U′_ varying from 1.1 to 1.5 %. This is not surprising as K ∼140 s^-1^ when *R*_2,U′_ = 200 s^-1^ resulting in relatively small *ω*_1_/K values of 0.27 and 0.82 for *B*_*1*_ fields of 6.0 and 18.4 Hz, respectively. For these seven residues with large *R*_2,U′_ values, significantly more precise exchange parameters were obtained when the CEST data recorded with a *B*_*1*_ of 46 Hz was also included in the analysis (Fig. 3h), with *k*_*ex*_ now varying from 57 to 90 s^-1^ and *p*_U′_ now varying from 1.2 to 1.5%. A *B*_*1*_ field of 46 Hz (*ω*_1_/*k*_*ex*_ ∼ 4) corresponds to a *ω*_1_/K value of 2.1 when *R*_2,U′_ = 200 s^-1^ and thus including this higher field results in more precise exchange rates when the minor state dips are severely broadened, which is consistent with the theoretical analysis presented above. In a previous study of the A39G FF folding using ^15^N CEST experiments precise exchange parameters were obtained because datasets with high *B*_*1*_ values were inadvertently recorded, while looking for the minor state dips (25).

**Fig. 3.**
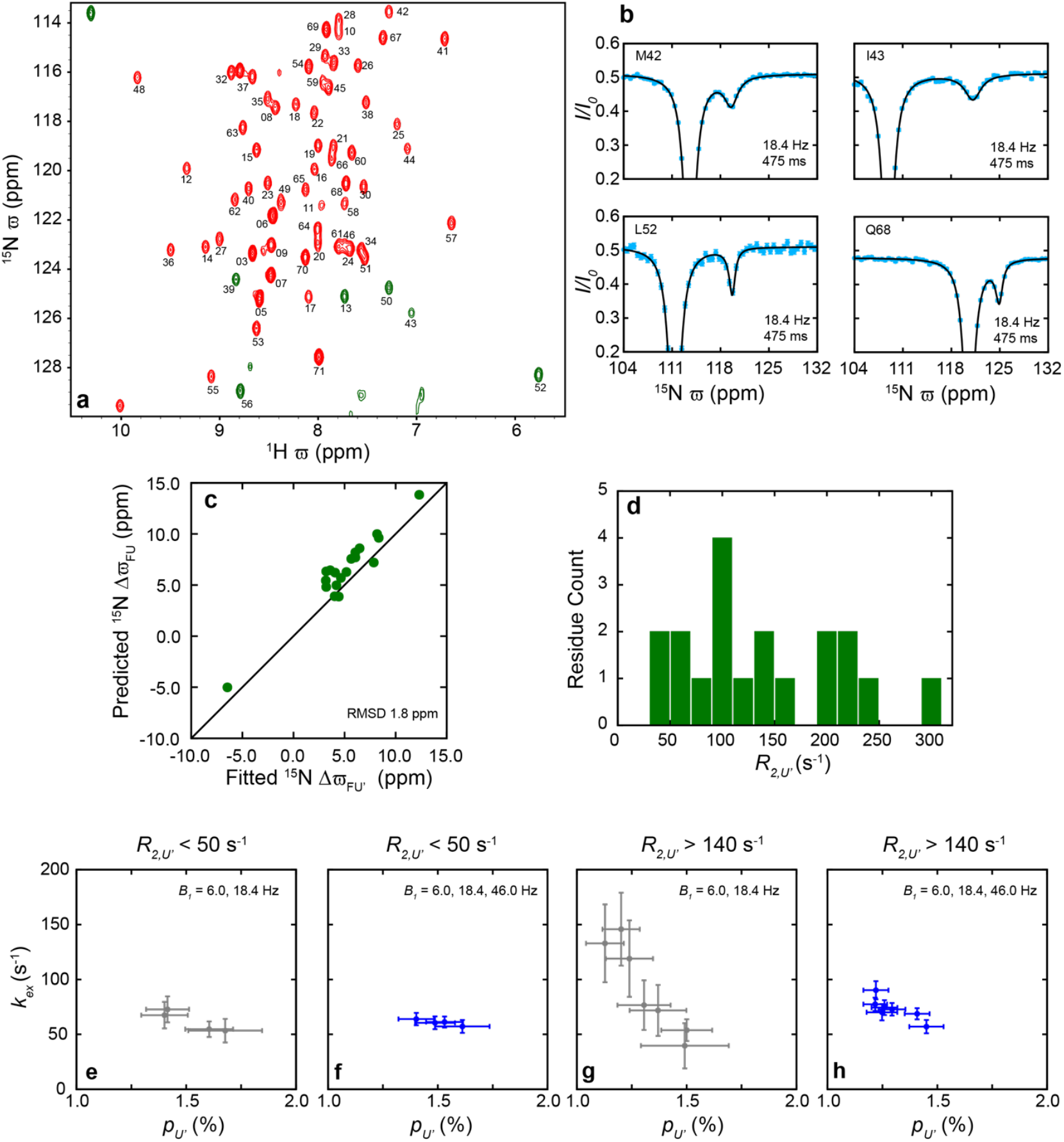
Folding of the A39G FF domain studied using ^15^N CEST experiments. (a) The amide ^15^N-^1^H correlation map of U-[^15^N] A39G FF (11.7 T, 2.5 °C) in which peaks are labelled according to the residue from which they arise. Peaks aliased in the ^15^N dimension are shown in green. (b) Representative amide ^15^N CEST profiles (*B*_*1*_ and *T*_*EX*_ indicated) from four different sites in the molecule. Cyan circles represent the experimental data and the black line is drawn according to global best fit parameters (*k*_*ex*_ = 71.6 s^-1^, *p*_*U*′_ = 1.39%; Table S3). (c) Correlation between the predicted 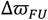 and CEST derived 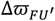 shifts. 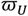 shifts were predicted using the program POTENCI (71). (d) Distribution of the *R*_2,*U*′_ values obtained from a global analysis 6.0 and 18.4 Hz ^15^N CEST data. (e) For the four residues with *R*_2,*U*′_ < 50 s^-1^ very similar residue specific *k*_*ex*_ and *p*_*U*′_ values are obtained from the analysis of 6.0 and 18.4 Hz ^15^N CEST data and the inclusion of 46.0 Hz CEST data does not really have an effect on the distribution of the *k*_*ex*_ and *p*_*U*′_ values (f). (g) For the seven residues with *R*_2,*U*′_ > 140 s^-1^ there is a large variation in the residue specific *k*_*ex*_ values obtained from the analysis 6.0 and 18.4 Hz ^15^N CEST data and the inclusion of 46.0 Hz CEST data leads to a narrower distribution of *k*_*ex*_ and *p*_*U*′_ values (h).

### Concluding remarks

We have shown that the choice of *B*_*1*_ fields required to characterize chemical exchange using CEST experiments depends on the (apparent) minor state transverse relaxation rate in addition to *k*_*ex*_. We suggest that the choice of *B*_*1*_ fields should be governed by 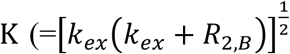 as opposed to *k*_*ex*_. When *R*_2,*B*_ ≪ *k*_*ex*_, K ≈ *k*_*ex*_ and the choice of *B*_*1*_ fields to characterize exchange will be essentially determined by *k*_*ex*_. However, when *R*_2,*B*_ is substantially greater than *k*_*ex*_, CEST datasets recorded with higher *B*_*1*_ fields determined by K, as opposed to *k*_*ex*_, are required to obtain accurate exchange parameters. Often this will necessitate recording an additional CEST dataset with a relatively high *B*_*1*_ value (recommended to be ∼4.5*k*_*ex*_/2*π*) so that 2*πB*_1_/K is greater than ∼1.8 for all the sites in the molecule. Although this strategy often requires recording additional CEST datasets with higher *B*_*1*_ values, it should be noted that these datasets can be recorded rapidly compared to datasets with lower *B*_*1*_ values as the spacing between adjacent offsets, at which *B*_*1*_ irradiation is carried out, is larger when the *B*_*1*_ values are higher (14, 70). We expect that the conclusions presented here will be valuable when CEST experiments are used to characterize slow processes (*k*_*ex*_ ≤ ∼25 s^-1^) in large proteins, processes with *k*_*ex*_ ≤ ∼10 s^-1^ in small to medium sized proteins and when the minor state dips are severely exchange broadened due to the presence of other sparsely populated states, as in the case of A39G FF studied here. These results will continue to become more relevant as higher field spectrometers become available because the transverse relaxation rates for several sites in protein molecules will increase with field strength.

## Supporting Information

Supporting Information is included in this file after the references.

## Acknowledgements

The authors thank Nemika Thapliyal (TIFR Hyderabad) and Dr Debajyoti De (TIFR Hyderabad) for providing the A17G S56P FF sample, Dr. G. Bouvignies (ENS, Paris) for providing the program *ChemEx*, Prof. Lewis Kay (University of Toronto) for providing the CEST and D-CEST pulse sequences, TIFR Hyderabad NMR facility for the generous grant of spectrometer time and Dr Krishna Rao for maintaining the facility. PV acknowledges intramural funding from TIFR Hyderabad (DAE, Government of India, Project No. RTI 4007). DFH is supported by the UKRI and EPSRC (EP/X036782/1). For the purpose of open access, the author has applied a Creative Commons Attribution (CC BY) license to any Author Accepted Manuscript version arising.

## Supporting Information

### Supporting Text

#### Calculations confirm that CEST datasets with ‘high’ *B*_*1*_ fields are necessary to obtain precise exchange parameters for slow processes when *R*_*2,B*_ is larger than *K*_*ex*_

Monte Carlo simulations (1, 2) were used to test the validity of the theoretical analysis presented in the text. CEST profiles (^15^N; 16.4 T) with *B*_*1*_ values of 1.7, 4.1, 9.9 and 11.7 Hz were generated for two ‘residues’ with *k*_***ex***_ = 15 s^-1^, *p*_*B*_ = 7.5 % and 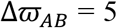 ppm. When *k*_***ex***_ is 15 s^-1^, *B*_*1*_ values of 1.7, 4.1, 9.9 and 11.7 Hz correspond to *ω*_1_/*k*_***ex***_ values of 0.7, 1.7, 4.1 and 4.9 respectively.

For residue 1 *R*_2,*B*_ was set to 5 s^-1^ resulting in K 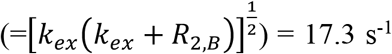 that is similar to the *k*_***ex***_ value of 15 s^-1^. Fits to the 1.7 and 4.1 Hz CEST profiles results in well-defined exchange parameters (*k*_***ex***_ = 15 ± 1.4 s^-1^, *p*_*B*_ = 7.5 ± 0.4 %; grey circles in Fig. S2a) and a distinct minimum in the 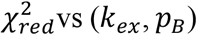 plot (Fig. S2b) because *B*_*1*_ values of 1.7 and 4.1 Hz correspond to *ω*_1_/K values of 0.6 and 1.5 respectively. Including the CEST profile calculated with *B*_*1*_= 11.7 Hz only has a small effect on the extracted exchange parameters (*k*_***ex***_ = 15 ± 0.7 s^-1^, *p*_*B*_ = 7.5 ± 0.3 %; blue pluses in Fig. S2a; Fig. S2c).

For residue 2 on the other hand, *R*_2,*B*_ was set to 75 s^-1^ resulting in K = 36.7 s^-1^ that is more than twice *k*_***ex***_ and fits to the 1.7 and 4.1 Hz CEST profiles results in poorly defined exchange parameters (*k*_***ex***_ = 15 ± 7 s^-1^, *p*_*B*_ =7.5 ± 1.0 %; grey circles in Fig. S2d) and a 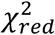 vs (*k*_***ex***_, *p*_*B*_) plot without a sharp minimum (especially along *k*_***ex***_) (Fig. S2e) because 1.7 and 4.1 Hz correspond to *ω*^*^/K values of 0.3 and 0.7 that are too small for the extraction of accurate exchange parameters. Including the CEST dataset calculated with *B*_*1*_ = 11.7 Hz in the analysis procedure leads to more precise exchange parameters (*k*_***ex***_ = 15 ± 0.8 s^-1^, *p*_*B*_ = 7.5 ± 0.3 %; blue pluses in Fig. S2d) and a distinct minimum in the 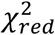 vs (*k*_***ex***_, *p*_*B*_) plot (Fig. S2f) because *B*_*1*_ = 11.7 Hz corresponds to a *ω*_1_/K value of 2 for residue 2 which nicely complements the *B*_*1*_ = 1.7 and 4.1 Hz datasets that correspond to *ω*_1_/K values of 0.3 and 0.7 respectively. Finally precise exchange parameters (*k*_***ex***_ = 15 ± 1 s^-1^, *p*_*B*_ = 7.5 ± 0.3 %; Fig. S2g,h) were also obtained by analyzing the 4.1 & 9.9 Hz CEST datasets that correspond to the ‘recommended’ *ω*_1_/K values of 0.7 and 1.7 (3).

In the above analysis the CEST profiles were generated with no errors but an uncertainty of 0.5 % in the normalized intensities was assumed to carry out the Monte Carlo analysis.

**Fig. S1.**
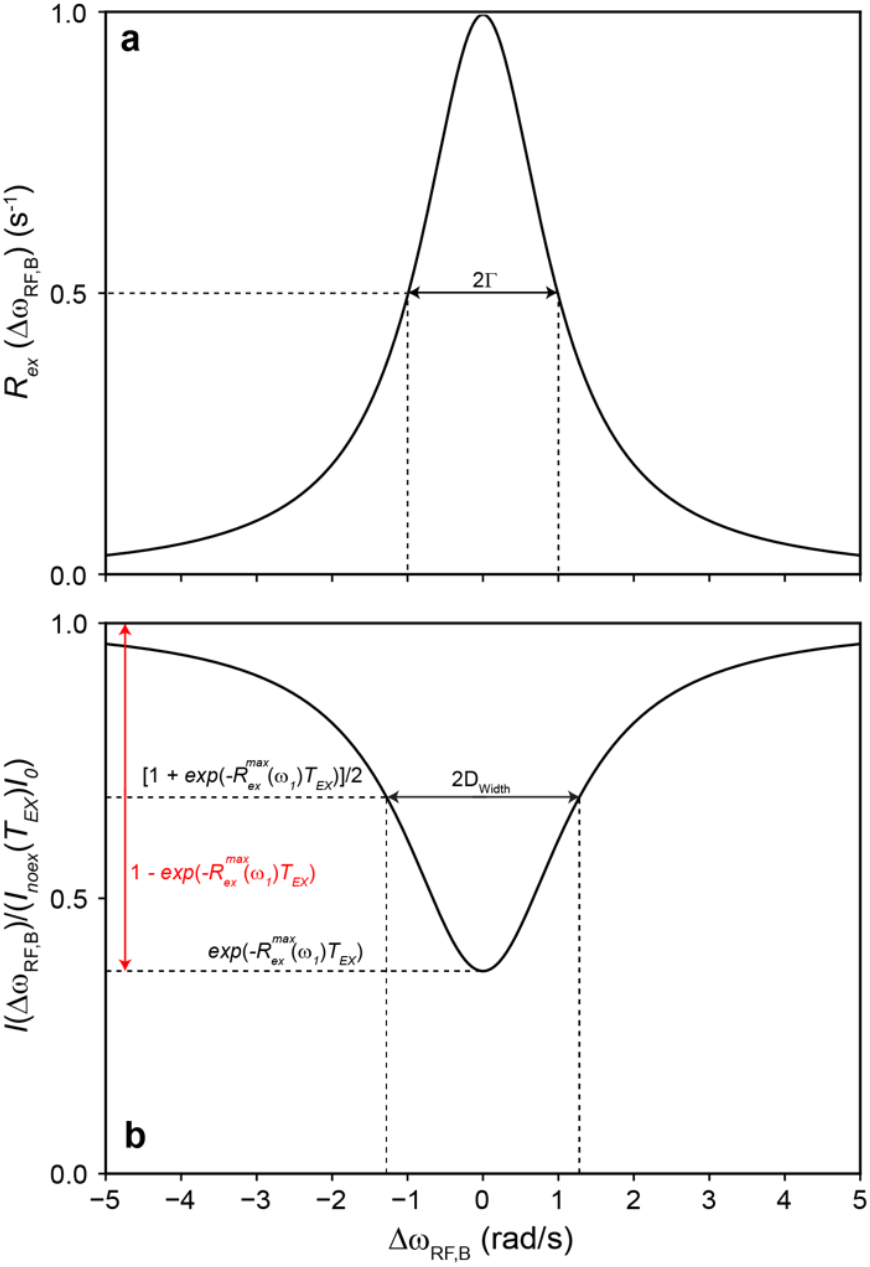
Schematic illustration of (a) *R*_***ex***_(Δ*ω*_*RF,B*_) 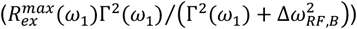 *v*.*s* Δ*ω*_*RF,B*_ and (b) 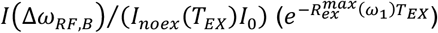 *v*.*s* Δ*ω*_*RF,B*_. According to equations 1-5 of the text (3-6), the size of the minor state dip is proportional to 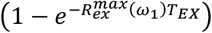 (red arrow in b) and its shape (*I*(Δ*ω*_*RF,B*_)/*I*_0_ *v*.*s* Δ*ω*_*RF,B*_) is proportional to 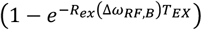. The “width” of the dip is 2D_width_ and D_width_ is the value of Δ*ω*_*RF,B*_ at which 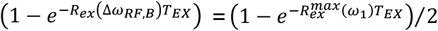. According to equation 2 in the text 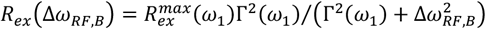 leading to 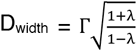, where 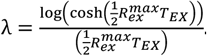. Hence D_width_ depends on Γ but is not Γ as Γ is the half width at half maximum of *R*_*ex*_(Δ*ω*_*RF,B*_). The plots were made with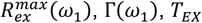 all set to 1 in their respective units.

**Fig. S2.**
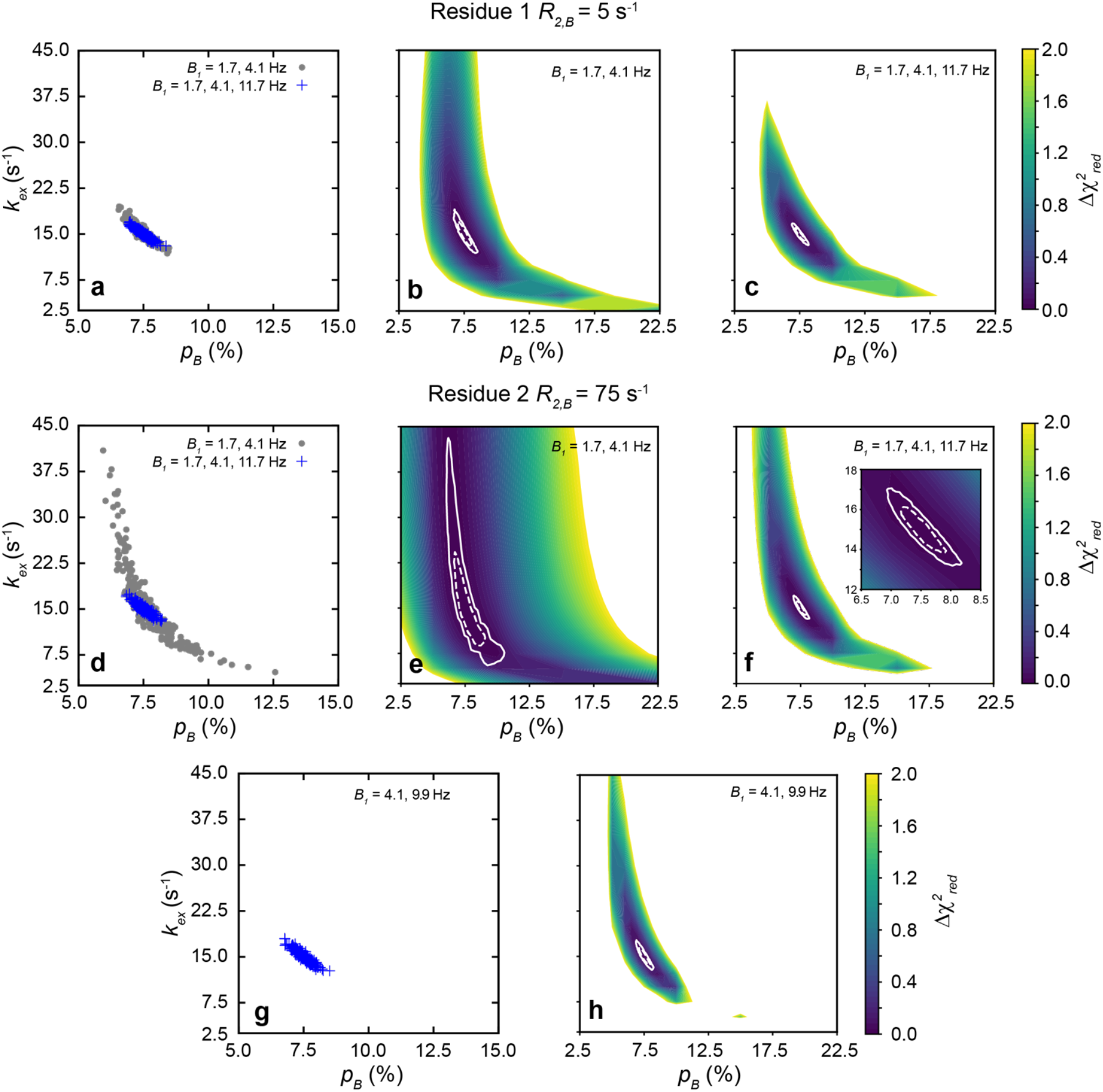
Simulations confirm that CEST datasets with ‘high’ *B*_*1*_ fields are necessary to obtain precise exchange parameters when *R*_2,*B*_ is high compared to *k*_***ex***_. Scatter plots of single residue exchange parameters obtained from a Monte Carlo procedure involving 250 trials carried out using calculated ^15^N CEST profiles (^15^N; 16.4 T; *T*_*EX*_ = 500 ms) that were generated for two “residues” with *k*_***ex***_ = 15 s^-1^, *p*_*B*_ = 7.5 %, *R*_1,*A*_ = *R*_1,*B*_ = 1 s^-1^, *R*_2,*A*_ = 5 s^-1^, 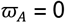 ppm, 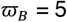 ppm, *R*_2,*B*_ = 5 s^-1^ for residue 1 (a) and *R*_2,*B*_ = 75 s^-1^ for residue 2 (d,g). 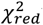 vs (*k*_***ex***_, *p*_*B*_) plots calculated for residue 1 (b,c) and residue 2 (e,f,h) by analyzing CEST datasets calculated using the indicated *B*_*1*_ values. It is clear that when *R*_2,*B*_ is high compared to *k*_***ex***_ (residue 2, panels d-h) that CEST datasets recorded with higher *B*_*1*_ values (9.9 or 11.7 Hz) are crucial for obtaining precise exchange parameters. In b,c,e,f and h 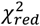 values above 2 are in white and contours corresponding to the 68 and 95% confidence intervals of *k*_***ex***_ and *p*_*B*_ based on 10,000 Monte Carlo trials are shown using dashed and solid white lines respectively. Here 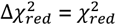 because the best fit 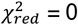 as the CEST profiles were generated with no errors.

**Table S1.**
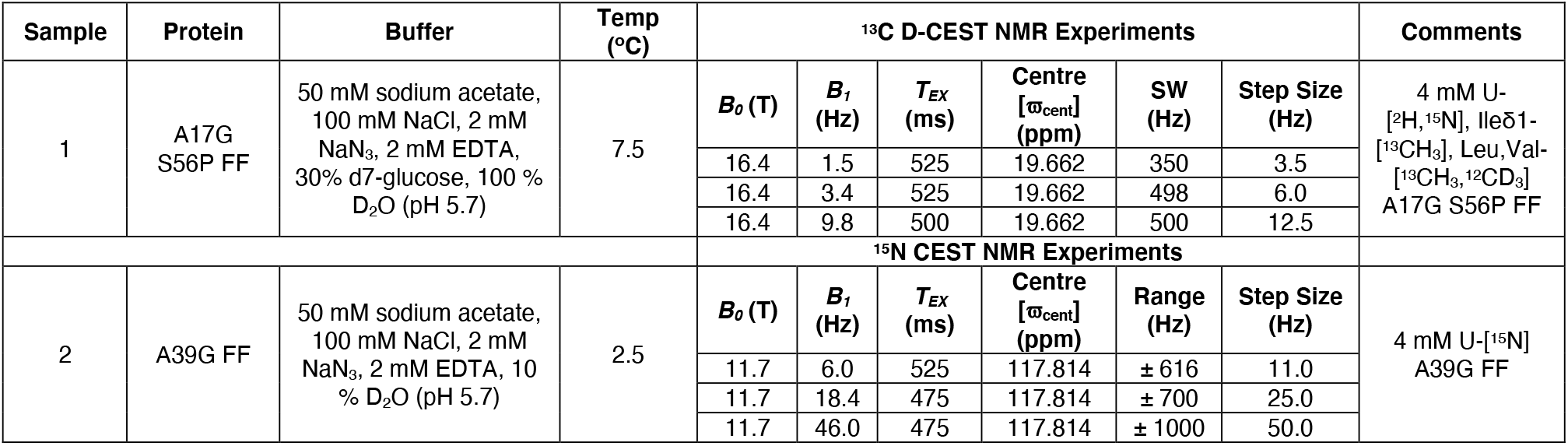
Details of the methyl ^13^C D-CEST (7) and the amide ^15^N CEST (8) experiments carried out in this study. SW is the sweep width of D-CEST sequence in the CEST dimension. *B*_*1*_ irradiation was carried out at offsets varying from -SW/2 (-Range) to +SW/2 (+Range) around ϖ_cent_ in steps of ‘Step Size’.

**Table S2.** Best fit exchange parameters obtained by analyzing two different sets of methyl ^13^C D-CEST profiles recorded using the 4 mM U-[^2^H,^15^N], Ileδ1-[^13^CH_3_], Leu,Val-[^13^CH_3_,^12^CD_3_] A17G S56P FF sample at 7.5 °C (16.4 T).

| Residue | $B_{1s}$<br>(Hz) | Global Analysis | | | | | | | | Residue-Specific Analysis | | | | | | | |
| --- | --- | --- | --- | --- | --- | --- | --- | --- | --- | --- | --- | --- | --- | --- | --- | --- | --- |
| | | $k_{ex}$ (s<sup>-1</sup>) | $p_I$ (%) | $\Delta\omega$ (ppm) | $R_{1,F}$ (s<sup>-1</sup>) | $R_{2,F}$ (s<sup>-1</sup>) | $R_{2,I}$ (s<sup>-1</sup>) | $\chi^2_{red}$ | K<br>(s<sup>-1</sup>) | $k_{ex}$ (s<sup>-1</sup>) | $p_I$ (%) | $\Delta\omega$ (ppm) | $R_{1,F}$ (s<sup>-1</sup>) | $R_{2,F}$ (s<sup>-1</sup>) | $R_{2,I}$ (s<sup>-1</sup>) | $\chi^2_{red}$ | K<br>(s<sup>-1</sup>) |
| V30<sub>γ</sub>2 | 1.5,<br>3.4 Hz | 15.9 ± 2.3 | 8.00 ± 0.42 | -0.30 ± 0.01 | 1.35 ± 0.01 | 17.2 ± 0.4 | 26.8 ± 2.9 | 0.93 | 26 | 6.4 ± 6.8 | 12.86 ± 9.45 | -0.30 ± 0.01 | 1.35 ± 0.01 | 18.1 ± 1.1 | 34.5 ± 7.2 | 1.01 | 16 |
| I43δ1 |  |  |  | 0.71 ± 0.01 | 0.86 ± 0.01 | 15.5 ± 0.4 | 24.6 ± 2.2 |  | 25 | 32.5 ± 9.3 | 6.82 ± 0.54 | 0.71 ± 0.01 | 0.86 ± 0.01 | 14.3 ± 0.7 | 13.6 ± 8.0 | 0.96 | 39 |
| I44δ1 |  |  |  | -0.29 ± 0.01 | 0.72 ± 0.01 | 11.8 ± 0.5 | 15.0 ± 2.9 |  | 22 | 12.1 ± 3.0 | 9.21 ± 0.95 | -0.28 ± 0.01 | 0.72 ± 0.01 | 12.1 ± 0.5 | 18.0 ± 2.3 | 0.51 | 19 |
| L52δ2 |  |  |  | 0.58 ± 0.01 | 1.30 ± 0.01 | 16.5 ± 0.7 | 14.1 ± 4.6 |  | 22 | 33.0 ± 5.7 | 6.25 ± 0.51 | 0.59 ± 0.01 | 1.31 ± 0.01 | 15.2 ± 0.6 | 0.0 ± 4.9 | 1.59 | 33 |
| L55δ1 |  |  |  | 0.86 ± 0.01 | 1.86 ± 0.01 | 13.6 ± 0.9 | 31.2 ± 9.5 |  | 27 | 17.3 ± 13.7 | 7.06 ± 1.92 | 0.86 ± 0.01 | 1.86 ± 0.01 | 13.3 ± 1.1 | 25.3 ± 10.7 | 0.77 | 27 |
| L55δ2 |  |  |  | 0.50 ± 0.01 | 2.41 ± 0.01 | 16.1 ± 0.8 | 50.3 ± 5.5 |  | 32 | 51.7 ± 18.9 | 5.37 ± 4.19 | 0.51 ± 0.01 | 2.42 ± 0.01 | 13.9 ± 1.5 | 7.8 ± 16.4 | 0.64 | 55 |
| Residue | $B_{1s}$<br>(Hz) | Global Analysis | | | | | | | | Residue-Specific Analysis | | | | | | | |
| | | $k_{ex}$ (s<sup>-1</sup>) | $p_I$ (%) | $\Delta\omega$ (ppm) | $R_{1,F}$ (s<sup>-1</sup>) | $R_{2,F}$ (s<sup>-1</sup>) | $R_{2,I}$ (s<sup>-1</sup>) | $\chi^2_{red}$ | K<br>(s<sup>-1</sup>) | $k_{ex}$ (s<sup>-1</sup>) | $p_I$ (%) | $\Delta\omega$ (ppm) | $R_{1,F}$ (s<sup>-1</sup>) | $R_{2,F}$ (s<sup>-1</sup>) | $R_{2,I}$ (s<sup>-1</sup>) | $\chi^2_{red}$ | K<br>(s<sup>-1</sup>) |
| V30<sub>γ</sub>2 | 1.5,<br>3.4,<br>9.8 Hz | 11.2 ± 0.5 | 9.52 ± 0.27 | -0.31 ± 0.01 | 1.36 ± 0.01 | 19.3 ± 0.3 | 35.5 ± 1.9 | 1.00 | 23 | 10.1 ± 1.9 | 10.68 ± 1.21 | -0.31 ± 0.01 | 1.36 ± 0.01 | 19.2 ± 0.6 | 36.7 ± 4.9 | 1.08 | 22 |
| I43δ1 |  |  |  | 0.71 ± 0.01 | 0.87 ± 0.01 | 17.1 ± 0.3 | 31.6 ± 1.6 |  | 22 | 10.4 ± 0.9 | 9.80 ± 0.52 | 0.71 ± 0.01 | 0.87 ± 0.01 | 17.2 ± 0.3 | 32.0 ± 2.0 | 0.97 | 21 |
| I44δ1 |  |  |  | -0.28 ± 0.01 | 0.72 ± 0.01 | 13.2 ± 0.3 | 22.0 ± 2.1 |  | 19 | 12.3 ± 1.0 | 9.64 ± 0.50 | -0.28 ± 0.01 | 0.72 ± 0.01 | 12.8 ± 0.3 | 22.5 ± 1.6 | 0.61 | 21 |
| L52δ2 |  |  |  | 0.58 ± 0.01 | 1.31 ± 0.01 | 18.0 ± 0.4 | 23.3 ± 3.0 |  | 20 | 12.4 ± 1.3 | 9.05 ± 0.60 | 0.59 ± 0.01 | 1.31 ± 0.01 | 17.8 ± 0.4 | 22.7 ± 2.1 | 1.66 | 21 |
| L55δ1 |  |  |  | 0.86 ± 0.01 | 1.86 ± 0.01 | 15.3 ± 0.6 | 32.8 ± 7.9 |  | 22 | 11.5 ± 1.7 | 8.48 ± 0.85 | 0.86 ± 0.01 | 1.87 ± 0.01 | 15.1 ± 0.5 | 33.5 ± 3.7 | 0.90 | 23 |
| L55δ2 |  |  |  | 0.50 ± 0.01 | 2.41 ± 0.01 | 16.6 ± 0.5 | 66.7 ± 4.4 |  | 30 | 11.7 ± 2.7 | 8.87 ± 1.18 | 0.50 ± 0.01 | 2.41 ± 0.01 | 16.7 ± 0.6 | 64.7 ± 8.0 | 0.69 | 30 |

**Table S3.**
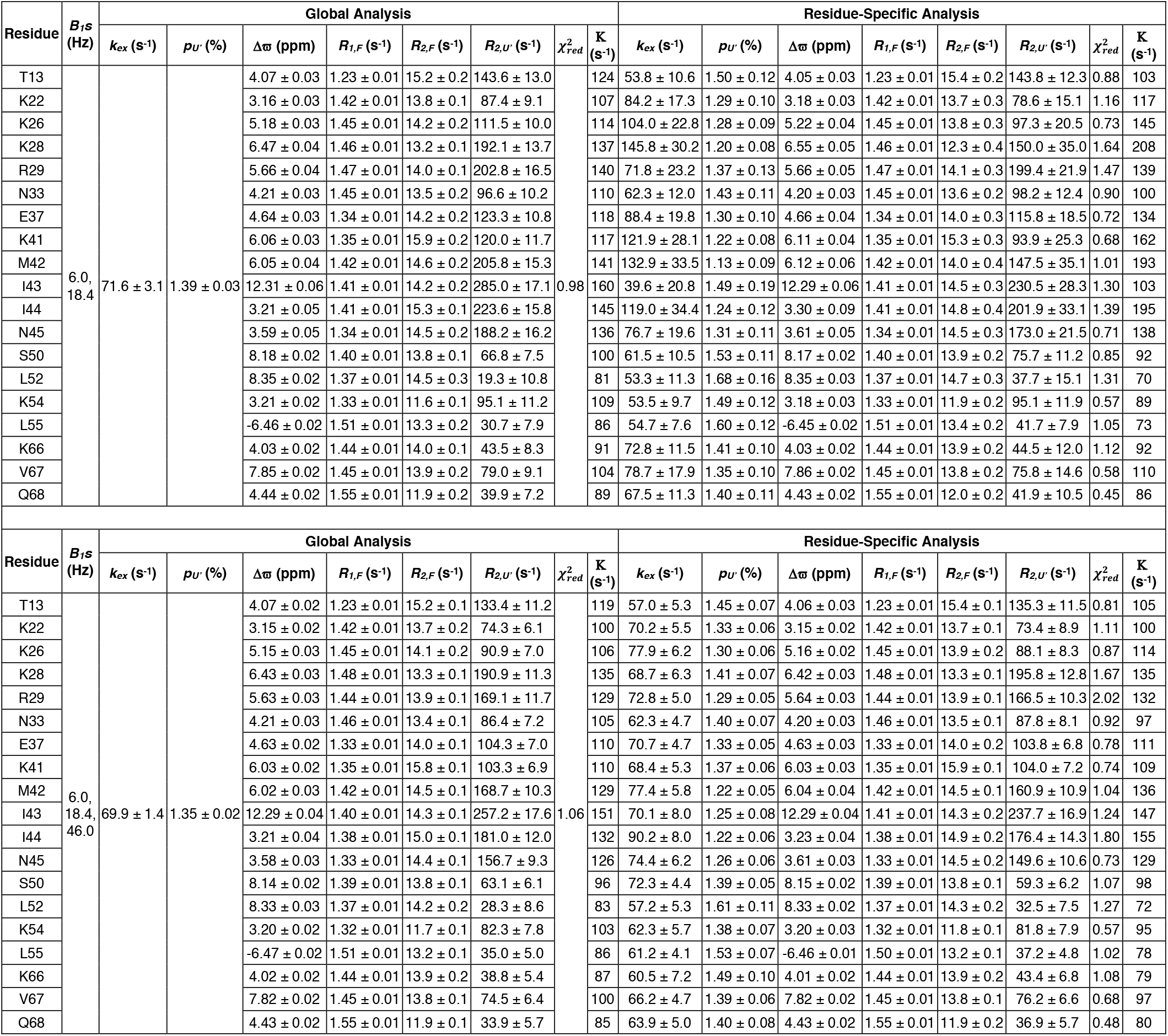
Best fit exchange parameters extracted from two different sets of ^15^N CEST profiles recorded using the 4 mM U-[^15^N] A39G FF sample at 2.5 ºC (11.7 T).

## References

1. Karplus M. Aspects of protein reaction dynamics: Deviations from simple behavior. J Phys Chem B. 2000;104(1):11–27.

2. Bahar I, Jernigan R, Dill KA. Protein actions : principles and modeling. New York: Garland Science, Taylor & Francis Group; 2017. xii, 322 pages p.

3. Karplus M, Kuriyan J. Molecular dynamics and protein function. Proc Natl Acad Sci U S A. 2005;102(19):6679–85.

4. Sekhar A, Kay LE. An NMR View of Protein Dynamics in Health and Disease. Annu Rev Biophys. 2019;48:297–319.

5. Lisi GP, Loria JP. Allostery in enzyme catalysis. Curr Opin Struct Biol. 2017;47:123–30.

6. Shukla VK, Siemons L, Hansen DF. Intrinsic structural dynamics dictate enzymatic activity and inhibition. Proc Natl Acad Sci U S A. 2023;120(41):e2310910120.

7. Zhuravleva A, Korzhnev DM. Protein folding by NMR. Prog Nucl Magn Reson Spectrosc. 2017;100:52–77.

8. Palmer AG, 3rd, Koss H. Chemical Exchange. Methods Enzymol. 2019;615:177–236.

9. Anthis NJ, Clore GM. Visualizing transient dark states by NMR spectroscopy. Q Rev Biophys. 2015;48(1):35–116.

10. Palmer AG, Massi F. Characterization of the dynamics of biomacromolecules using rotating-frame spin relaxation NMR spectroscopy. Chem Rev. 2006;106(5):1700–19.

11. Rangadurai A, Szymaski ES, Kimsey IJ, Shi H, Al-Hashimi H. Characterizing micro-to-millisecond chemical exchange in nucleic acids using off-resonance R1ρ relaxation dispersion. Progress in Nuclear Magnetic Resonance Spectroscopy. 2019;112-113:55–102.

12. Palmer AG, 3rd, Kroenke CD, Loria JP. Nuclear magnetic resonance methods for quantifying microsecond-to-millisecond motions in biological macromolecules. Methods Enzymol. 2001;339:204–38.

13. Sauerwein A, Hansen DF. Relaxation Dispersion NMR Spectroscopy. In: Berliner L, editor. Protein NMR Biological Magnetic Resonance and Biomedical Applications. 32. Boston, MA: Springer; 2015. p. 75–132.

14. Vallurupalli P, Sekhar A, Yuwen T, Kay LE. Probing conformational dynamics in biomolecules via chemical exchange saturation transfer: a primer. J Biomol NMR. 2017;67(4):243–71.

15. Tugarinov V, Clore GM. Exchange saturation transfer and associated NMR techniques for studies of protein interactions involving high-molecular-weight systems. J Biomol NMR. 2019;73(8-9):461–9.

16. Bouvignies G, Vallurupalli P, Hansen DF, Correia BE, Lange O, Bah A, et al. Solution structure of a minor and transiently formed state of a T4 lysozyme mutant. Nature. 2011;477(7362):111–4.

17. Vallurupalli P, Hansen DF, Kay LE. Structures of invisible, excited protein states by relaxation dispersion NMR spectroscopy. Proc Natl Acad Sci U S A. 2008;105(33):11766–71.

18. Kukic P, Pustovalova Y, Camilloni C, Gianni S, Korzhnev DM, Vendruscolo M. Structural Characterization of the Early Events in the Nucleation-Condensation Mechanism in a Protein Folding Process. J Am Chem Soc. 2017;139(20):6899–910.

19. Neudecker P, Robustelli P, Cavalli A, Walsh P, Lundstrom P, Zarrine-Afsar A, et al. Structure of an intermediate state in protein folding and aggregation. Science. 2012;336(6079):362–6.

20. Hansen DF, Vallurupalli P, Kay LE. Using relaxation dispersion NMR spectroscopy to determine structures of excited, invisible protein states. J Biomol NMR. 2008;41(3):113–20.

21. Forsen S, Hoffman RA. Study of Moderately Rapid Chemical Exchange Reactions by Means of Nuclear Magnetic Double Resonance. J Chem Phys. 1963;39(11):2892–901.

22. Rangadurai A, Shi H, Al-Hashimi HM. Extending the Sensitivity of CEST NMR Spectroscopy to Micro-to-Millisecond Dynamics in Nucleic Acids Using High-Power Radio-Frequency Fields. Angew Chem Int Ed Engl. 2020;59(28):11262–6.

23. Khandave NP, Sekhar A, Vallurupalli P. Studying micro to millisecond protein dynamics using simple amide (15)N CEST experiments supplemented with major-state R(2) and visible peak-position constraints. J Biomol NMR. 2023.

24. Tiwari VP, Toyama Y, De D, Kay LE, Vallurupalli P. The A39G FF domain folds on a volcano-shaped free energy surface via separate pathways. Proc Natl Acad Sci U S A. 2021;118(46).

25. Vallurupalli P, Bouvignies G, Kay LE. Studying “invisible” excited protein States in slow exchange with a major state conformation. J Am Chem Soc. 2012;134(19):8148–61.

26. Tiwari VP, Vallurupalli P. A CEST NMR experiment to obtain glycine (1)H(alpha) chemical shifts in‘invisible’ minor states of proteins. J Biomol NMR. 2020;74(8-9):443–55.

27. Vallurupalli P, Kay LE. Probing slow chemical exchange at carbonyl sites in proteins by chemical exchange saturation transfer NMR spectroscopy. Angew Chem Int Ed Engl. 2013;52(15):4156–9.

28. Karunanithy G, Reinstein J, Hansen DF. Multiquantum Chemical Exchange Saturation Transfer NMR to Quantify Symmetrical Exchange: Application to Rotational Dynamics of the Guanidinium Group in Arginine Side Chains. J Phys Chem Lett. 2020;11(14):5649–54.

29. Yuwen T, Sekhar A, Kay LE. Separating dipolar and chemical exchange magnetization transfer processes in 1H-CEST. Angew Chem Int Ed Engl. 2017;56(22):6122–5.

30. Vallurupalli P, Bouvignies G, Kay LE. A Computational Study of the Effects of C-13-C-13 Scalar Couplings on C-13 CEST NMR Spectra: Towards Studies on a Uniformly C-13-Labeled Protein. Chembiochem. 2013;14(14):1709–13.

31. Bouvignies G, Vallurupalli P, Kay LE. Visualizing Side Chains of Invisible Protein Conformers by Solution NMR. Journal of Molecular Biology. 2014;426(3):763–74.

32. Vallurupalli P, Tiwari VP, Ghosh S. A Double-Resonance CEST Experiment To Study Multistate Protein Conformational Exchange: An Application to Protein Folding. J Phys Chem Lett. 2019;10(11):3051–6.

33. Madhurima K, Nandi B, Munshi S, Naganathan AN, Sekhar A. Functional regulation of an intrinsically disordered protein via a conformationally excited state. Sci Adv. 2023;9(26):eadh4591.

34. Gladkova C, Schubert AF, Wagstaff JL, Pruneda JN, Freund SMV, Komander D. An invisible ubiquitin conformation is required for efficient phosphorylation by PINK1. EMBO J. 2017;36(24):3555–72.

35. Zhao B, Hansen AL, Zhang Q. Characterizing Slow Chemical Exchange in Nucleic Acids by Carbon CEST and Low Spin-Lock Field R1ρ NMR Spectroscopy. J Am Chem Soc. 2014;136(1):20–3.

36. Lim J, Xiao TS, Fan JS, Yang DW. An Off-Pathway Folding Intermediate of an Acyl Carrier Protein Domain Coexists with the Folded and Unfolded States under Native Conditions. Angew Chem Int Ed Engl. 2014;53(9):2358–61.

37. Tiwari VP, D. D, Thapliyal N, Kay LE, Vallurupalli P. Beyond slow two-state protein conformational exchange using CEST: applications to three-state protein interconversion on the millisecond timescale. J Biomol NMR. 2024.

38. Allen M, Friedler A, Schon O, Bycroft M. The structure of an FF domain from human HYPA/FBP11. Journal of Molecular Biology. 2002;323(3):411–6.

39. Korzhnev DM, Religa TL, Banachewicz W, Fersht AR, Kay LE. A transient and low-populated protein-folding intermediate at atomic resolution. Science. 2010;329(5997):1312–6.

40. Jemth P, Johnson CM, Gianni S, Fersht AR. Demonstration by burst-phase analysis of a robust folding intermediate in the FF domain. Protein Eng Des Sel. 2008;21(3):207–14.

41. Jemth P, Gianni S, Day R, Li B, Johnson CM, Daggett V, et al. Demonstration of a low-energy on-pathway intermediate in a fast-folding protein by kinetics, protein engineering, and simulation. Proc Natl Acad Sci U S A. 2004;101(17):6450–5.

42. Korzhnev DM, Religa TL, Lundstrom P, Fersht AR, Kay LE. The folding pathway of an FF domain: characterization of an on-pathway intermediate state under folding conditions by (15)N, (13)C(alpha) and (13)C-methyl relaxation dispersion and (1)H/(2)H-exchange NMR spectroscopy. J Mol Biol. 2007;372(2):497–512.

43. Jemth P, Day R, Gianni S, Khan F, Allen M, Daggett V, et al. The structure of the major transition state for folding of an FF domain from experiment and simulation. J Mol Biol. 2005;350(2):363–78.

44. Korzhnev DM, Vernon RM, Religa TL, Hansen AL, Baker D, Fersht AR, et al. Nonnative interactions in the FF domain folding pathway from an atomic resolution structure of a sparsely populated intermediate: an NMR relaxation dispersion study. J Am Chem Soc. 2011;133(28):10974–82.

45. Goto NK, Gardner KH, Mueller GA, Willis RC, Kay LE. A robust and cost-effective method for the production of Val, Leu, Ile (delta 1) methyl-protonated 15N-, 13C-, 2H-labeled proteins. J Biomol NMR. 1999;13(4):369–74.

46. Tugarinov V, Kay LE. Methyl groups as probes of structure and dynamics in NMR studies of high-molecular-weight proteins. Chembiochem. 2005;6(9):1567–77.

47. Gopalan AB, Vallurupalli P. Measuring the signs of the methyl 1H chemical shift diferences between major and‘invisible’ minor protein conformational states using methyl 1H multi-quantum spectroscopy. J Biomol NMR. 2018;70(3):187–202.

48. Vallurupalli P, Hansen DF, Lundstrom P, Kay LE. CPMG relaxation dispersion NMR experiments measuring glycine 1H alpha and 13C alpha chemical shifts in the‘invisible’ excited states of proteins. J Biomol NMR. 2009;45(1-2):45–55.

49. Yuwen T, Bouvignies G, Kay LE. Exploring methods to expedite the recording of CEST datasets using selective pulse excitation. J Magn Reson. 2018;292:1–7.

50. Yuwen T, Kay LE, Bouvignies G. Dramatic Decrease in CEST Measurement Times Using MultiSite Excitation. Chemphyschem. 2018;19(14):1707–10.

51. Bodenhausen G, Freeman R, Morris GA. Simple Pulse Sequence for Selective Excitation in Fourier-Transform Nmr. J Magn Reson. 1976;23(1):171–5.

52. Morris GA, Freeman R. Selective Excitation in Fourier-Transform Nuclear Magnetic-Resonance. J Magn Reson. 1978;29(3):433–62.

53. Levitt MH. Symmetrical Composite Pulse Sequences for Nmr Population-Inversion .2. Compensation of Resonance Offset. J Magn Reson. 1982;50(1):95–110.

54. Guenneugues M, Berthault P, Desvaux H. A method for determining B1 field inhomogeneity. Are the biases assumed in heteronuclear relaxation experiments usually underestimated? J Magn Reson. 1999;136(1):118–26.

55. Delaglio F, Grzesiek S, Vuister GW, Zhu G, Pfeifer J, Bax A. NMRPipe - a Multidimensional Spectral Processing System Based on Unix Pipes. J Biomol NMR. 1995;6(3):277–93.

56. Lee W, Tonelli M, Markley JL. NMRFAM-SPARKY: enhanced software for biomolecular NMR spectroscopy. Bioinformatics. 2015;31(8):1325–7.

57. Goddard TD, Kneller DG. SPARKY 3 University of California, San Francisco2008.

58. Ahlner A, Carlsson M, Jonsson BH, Lundstrom P. PINT: a software for integration of peak volumes and extraction of relaxation rates. J Biomol NMR. 2013;56(3):191–202.

59. Bouvignies G. Chemex (https://github.com/gbouvignies/chemex/releases)2012.

60. Korzhnev DM, Salvatella X, Vendruscolo M, Di Nardo AA, Davidson AR, Dobson CM, et al. Low-populated folding intermediates of Fyn SH3 characterized by relaxation dispersion NMR. Nature. 2004;430(6999):586–90.

61. McConnell HM. Reaction Rates by Nuclear Magnetic Resonance. J Chem Phys. 1958;28(3):430–1.

62. Press WH, Flannery BP, Teukolsky SA, Vetterling WT. Numerical Recipes in C. The Art of Scientific Computing Second Edition ed .Cambridge (UK): Cambridge University Press; 1992.

63. Choy WY, Zhou Z, Bai Y, Kay LE. An 15N NMR spin relaxation dispersion study of the folding of a pair of engineered mutants of apocytochrome b562. J Am Chem Soc. 2005;127(14):5066–72.

64. Mulder FA, Mittermaier A, Hon B, Dahlquist FW, Kay LE. Studying excited states of proteins by NMR spectroscopy. Nat Struct Biol. 2001;8(11):932–5.

65. Zaiss M, Bachert P. Chemical exchange saturation transfer (CEST) and MR Z-spectroscopy in vivo: a review of theoretical approaches and methods. Phys Med Biol. 2013;58(22):R221–69.

66. Zaiss M, Schnurr M, Bachert P. Analytical solution for the depolarization of hyperpolarized nuclei by chemical exchange saturation transfer between free and encapsulated xenon (HyperCEST). J Chem Phys. 2012;136(14):144106.

67. Baldwin AJ, Kay LE. An R(1rho) expression for a spin in chemical exchange between two sites with unequal transverse relaxation rates. J Biomol NMR. 2013;55(2):211–8.

68. Trott O, Palmer AG, 3rd. R1rho relaxation outside of the fast-exchange limit. J Magn Reson. 2002;154(1):157–60.

69. Miloushev VZ, Palmer AG, 3rd. R(1rho) relaxation for two-site chemical exchange: general approximations and some exact solutions. J Magn Reson. 2005;177(2):221–7.

70. Bolik-Coulon N, Hansen DF, Kay LE. Optimizing frequency sampling in CEST experiments. J Biomol NMR. 2022;76(5-6):167–83.

71. Nielsen JT, Mulder FAA. POTENCI: prediction of temperature, neighbor and pH-corrected chemical shifts for intrinsically disordered proteins. J Biomol NMR. 2018;70(3):141–65.

## References

1. Choy WY, Zhou Z, Bai Y, Kay LE. An 15N NMR spin relaxation dispersion study of the folding of a pair of engineered mutants of apocytochrome b562. J Am Chem Soc. 2005;127(14):5066–72.

2. Press WH, Flannery BP, Teukolsky SA, Vetterling WT. Numerical Recipes in C. The Art of Scientific Computing Second Edition ed. Cambridge (UK): Cambridge University Press; 1992.

3. Vallurupalli P, Sekhar A, Yuwen T, Kay LE. Probing conformational dynamics in biomolecules via chemical exchange saturation transfer: a primer. J Biomol NMR. 2017;67(4):243–71.

4. Zaiss M, Bachert P. Chemical exchange saturation transfer (CEST) and MR Z-spectroscopy in vivo: a review of theoretical approaches and methods. Phys Med Biol. 2013;58(22):R221–69.

5. Zaiss M, Schnurr M, Bachert P. Analytical solution for the depolarization of hyperpolarized nuclei by chemical exchange saturation transfer between free and encapsulated xenon (HyperCEST). J Chem Phys. 2012;136(14):144106.

6. Palmer AG, 3rd, Koss H. Chemical Exchange. Methods Enzymol. 2019;615:177–236.

7. Yuwen T, Kay LE, Bouvignies G. Dramatic Decrease in CEST Measurement Times Using Multi-Site Excitation. Chemphyschem. 2018;19(14):1707–10.

8. Vallurupalli P, Bouvignies G, Kay LE. Studying “invisible” excited protein States in slow exchange with a major state conformation. J Am Chem Soc. 2012;134(19):8148–61.

